# Rare shell colours in bivalves reveal multiple evolutionary pathways to blue and green colouration

**DOI:** 10.64898/2026.01.21.700781

**Authors:** Abigail L. Ingram, Joseph Razzell-Hollis, Suzanne T. Williams

## Abstract

Green and blue are rare colours in bivalve shells. Where such colours occur, it is unclear whether they result from shared pigments across taxa, as similar appearances may arise from different compounds and are not necessarily homologous. Identifying their biochemical basis is therefore crucial to understanding the evolution and possible metabolic costs of these unusual traits. We used Raman spectroscopy to analyse resin-embedded cross-sections of valves from 15 bivalve species with green or blue shells. Species represented diversity in shell colour in terms of both phylogeny and morphological traits, including colour location (organic periostracum, organic layers in the calcareous shell or inorganic calcareous matrix). Green colour consistently occurred in organic layers rather than calcareous shell, while blue colour occurs in both calcareous shell and organic layers. Blue colour appears to be due to carotenoid-based pigments, similar to other pigments observed in bivalve shells. Green colour, however, is due either to novel pigments, not previously identified in mollusc shells, which are only weakly Raman active or nanostructures that produce structural colour in the absence of green pigments. Within bivalves, to date structural colour has been reported only in one genus, underscoring its evolutionary significance.

## Background

Green and blue colours are surprisingly rare in the animal kingdom, and pigments producing these colours are even rarer (1). The fact that some colours are rare might be explained because that particular colour serves no useful function, or because the colour is difficult to synthesise (2). Grant & Williams (3) suggest the latter may be the case in bivalve shells and that these colours may provide some selective advantage since they have arisen multiple times in independent lineages. Only a handful of blue or green pigments have been identified in Mollusca to date, and most have been from the Vetigastropoda (4–9). It is not known whether these pigments are shared across taxonomic groups, since similar shell colours can arise from different pigments and are not necessarily homologous (10) and most surveys have focussed on more common shell colours (e.g. red, orange, brown, yellow, purple, etc.) (8, 10–16).

Curiously, previous observational studies have shown that in bivalves, rare blue and green shell colours are more commonly associated with the periostracum, an organic layer overlying the calcareous shell, rather than the mineralised shell itself (3). This led to the suggestion that the characteristics of the molecular structure of some blue and green bivalve pigments may mean some are more easily incorporated into organic than inorganic material (3). Identifying these pigments and determining how they are produced would be the first step towards understanding how and why these unusual, and possibly metabolically costly, pigments have evolved. An alternate suggestion to explain the distribution of rare colours, arises from previous Raman studies. Raman spectroscopy is frequently used with success to identify shell pigments in molluscs (e.g. (4, 17)), but Wade et al. (17) showed that the green layers on the inside of three bivalve shells, which are likely comprised of a mixture of organic and inorganic material, are Raman inactive – meaning that the constituent molecules have no vibrational modes readily detectable to Raman spectroscopy (17). The finding by Wade and colleagues is the first report of Raman inactive colouration in molluscs. The authors suggested this finding may be the result of novel green pigment(s) with weak or no Raman activity or the presence of structural colour, or a combination of both. Conversely, previous chemical analyses of the outer inorganic, mineral shell layer of green nerite gastropods and turban snail shells have identified green pigments as polyenes and bilins respectively (4–8), both of which are Raman active and can be detected by Raman spectroscopy. The explanation for this disparity may be due to the long evolutionary divergence between bivalves and gastropods and as previously suggested, in the location of pigments: in three green bivalves tested by Wade and colleagues, colour occurred in organic material.

Another possible explanation for the apparent lack of pigment in green bivalve shells, is the existence of structural colour. This has rarely been identified in bivalves, apart from in two green species of mussel: *Perna viridis* (18) and *P. canaliculus* (19), both of which possess a photonic periostracal layer, producing green iridescence. We plan to address these gaps in our understanding by investigating the evolution of rare blue and green bivalve shell colours and testing three hypotheses raised in previous studies, across a taxonomically diverse group of species: 1) rare green or blue colours occur more often in organic shell materials (periostracum and organic layers in calcareous shell) than calcareous shell matrix; 2) green or blue pigments associated with organic shell materials are Raman-inactive; and 3) green or blue colours in organic shell materials are the result of structural colour. Here, as a first step towards addressing these hypotheses, we use Raman spectroscopy to test the first two hypotheses and identify candidate species for further investigation of the third hypothesis in separate studies. We focus on 15 bivalve species that exhibit rare green and blue shell colours. Species were chosen to reflect the diversity of shell structures that can exhibit blue or green colouration (organic periostracum or organic layers in calcareous shell vs calcareous shell matrix) and the phylogenetic diversity of these species.

## Methods

### Sample selection and preparation

Green and blue shells were selected from NHMUK Mollusca collections, other than *Monia zelandica*, which was collected for this study by Dr Kerry Walton (Te Papa Tongarewa, Wellington, Museum of New Zealand), and the blue morph of *Septifer bilocularis*, which was purchased. The families were chosen to represent major clades: subclass Protobranchia (*Yoldia limatula*), infraclass Pteriomorphia (*Septifer bilocularis* – blue and green morphs, *Alectryonella plicatula, Arcuatula arcuatula, Monia zelandica, Perna viridis, Musculus discors, Mytella guyanensis*), superorder Imparidentia (*Corbicula fluminea, Glauconome virens, Chama croceata*), subterclass Palaeoheterodonta (*Anodonta anatina, Mutela legumen*) and superorder Anomalodesmata (*Cleidothaerus albidus*) (Fig. 1). No green or blue species have been identified in the Archiheterodonta (3).

**Figure 1.**
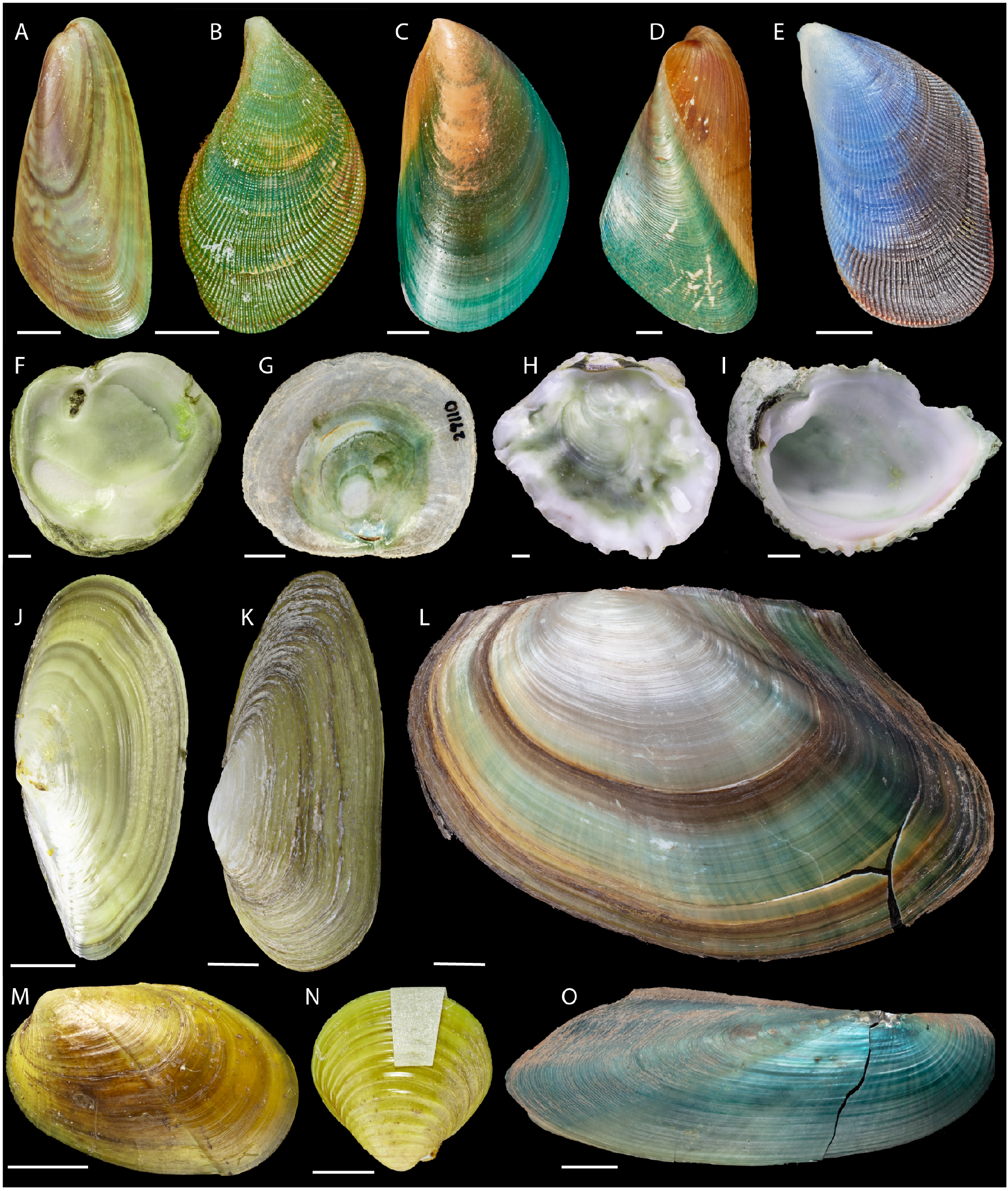
Photographs of 15 shells used in Raman analyses showing blue or green surfaces. All green and blue shell colour descriptions are based on Munsell Colour Charts (see Supporting Data, Figure 1). A, *Arcuatula arcuatula* (Hanley, 1844), NHMUK 20230896/1, showing ‘Yellow-Green-Yellow’ and brown striped periostracum on the outer valve. B, *Septifer bilocularis* (Linnaeus, 1758) green specimen NHMUK 1953-1-29 174-179, showing ‘Green-Yellow-Green’ periostracum on the outer valve. C, *Perna viridis* (Linnaeus, 1758) NHMUK 20230895_1, showing ‘bluish-Green’ periostracum on the outer valve. D, *Mytella guyanensis* (Lamarck, 1819) NHMUK 20230912, showing ‘Green’ and brown periostracum on the outer valve. E, *Septifer bilocularis* (Linnaeus, 1758) blue, showing ‘Purple-Blue’ and brown calcareous shell of the outer valve with periostracum removed. F, *Cleidothaerus albidus* (Lamarck, 1819), NHMUK 20230910, showing ‘greenish-Yellow’ inner valve. G, *Monia zelandica* (J. E. Gray, 1843), MusNZ029110_1, showing ‘greenish-Green-Yellow’ inner valve. H, *Alectryonella plicatula* (Gmelin, 1791), NHMUK 20230907/1, showing ‘greenish-Green-Yellow’ inner valve. I, *Chama croceata* Lamarck, 1819, NHMUK 1952.12.19 85, showing ‘greenish-Green-Yellow’ inner valve. J, *Yoldia limatula* (Say, 1831), NHMUK 20230858, showing ‘greenish-Yellow’ periostracum on the outer valve. K, *Glauconome virens* (Linnaeus, 1767), NHMUK 20240252/1, showing ‘greenish-Yellow’ periostracum on the outer valve. L, *Anodonta anatina* (Linnaeus, 1758), NHMUK 20180293, showing ‘Green’ and brown striped periostracum on the outer valve. M, *Musculus discors* (Linnaeus, 1767), NHMUK 77.11.28.15, showing ‘greenish-Yellow’ periostracum on the outer valve. N, *Corbicula fluminea* (O. F. Müller, 1774), NHMUK 20050005, showing ‘greenish-Yellow’ periostracum on the outer valve. O, *Mutela legumen* Graf & Cummings, 2006, NHMUK 1904.12.13-3-4, showing ‘greenish-Blue-Green’ periostracum on the outer valve. Scale bar: 5 mm. Images were taken with a QPcard 101 to allow for colour correction of final images.

Observers can differ in their opinion of colour; therefore, shell colours of specimens were determined by comparison with coloured charts in the Munsell Book of Color (20). Intact shell valves were photographed separately and alongside Munsell colour charts (Fig. 1 and Supporting Information Figure S1). Names used hereafter for shell colours refer to Munsell colours (Fig. 1 and S1).

Slide-mounted cross-sections of a single valve from each of the shells were prepared, enabling the use of light microscopy to provide a reference image and to precisely locate and investigate colour in the calcareous shell or the periostracum using Raman spectroscopy. Valves were embedded in KEPT epoxy resin (Kemet International Ltd, Maidstone, UK), before being cut in half from the centre of the umbo to the shell margin. The exposed surface was then polished to remove the rock saw marks using silicon carbide papers (2500 grit), reducing the thickness of the samples to 130 µm. Samples were then further polished using diamond and aluminium pastes of different grades to produce a 100 µm slide-mounted section. Colour images of shell cross-sections were taken in reflection on a white stage insert, using a Hirox HRX-01 digital light microscope (Fig. 2).

**Figure 2.**
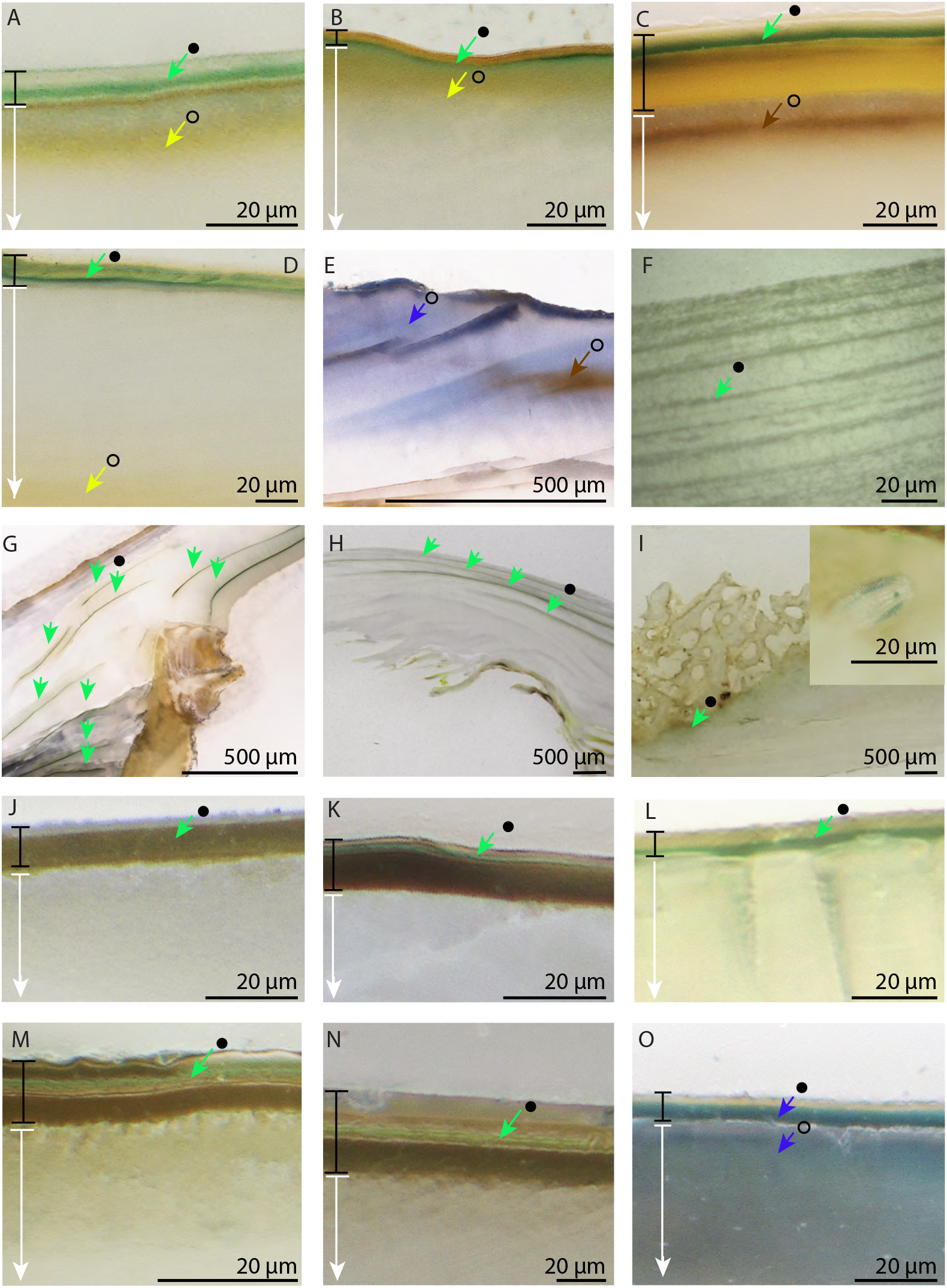
Photographs of resin-embedded cross-sections of shells, showing detail of green and blue colouration. A, *Arcuatula arcuatula*, showing three distinct periostracal layers: green middle layer flanked by two colourless layers, overlying diffuse yellow calcareous shell matrix. B, *Septifer bilocularis* (green), showing green basal and yellow upper periostracal layers, overlying diffuse yellow calcareous shell matrix. C, *Perna viridis*, showing three distinct periostracal layers: green middle layer flanked by colourless upper and yellow-brown basal layers, overlying brown calcareous shell matrix. D, *Mytella guyanensis*, showing three indistinct periostracal layers: uppermost yellow layer, green middle layer and colourless basal layer, overlying yellow calcareous shell matrix. E, *Septifer bilocularis* (blue), showing diffuse blue and brown colouration in the calcareous shell matrix. F, *Cleidothaerus albidus*, showing indistinct greenish yellow layers interleaved with white calcareous shell. G, *Monia zelandica*, showing green organic layers interleaved with white calcareous shell. H, *Alectryonella plicatula*, showing green organic layers interleaved with white calcareous shell. I, *Chama croceata*, showing diffuse green layer below the bioeroded shell, and above white calcareous shell; insert – green diatom in recesses of bioeroded shell. J, *Yoldia limatula*, showing two layers: outer colourless layer and olive-brown basal layer, overlying white calcareous shell. K, *Glauconome virens*, showing three periostracal layers: green middle layer, brown upper and basal layers, overlying white calcareous shell. L, *Anodonta anatina* showing two layers visible: outer colourless layer and green basal layer, overlying white calcareous shell. M, *Musculus discors*, showing three distinct periostracal layers: green middle layer, brown upper and basal layers, overlying white calcareous shell. N, *Corbicula fluminea*, showing three distinct periostracal layers: green middle layer, light brown upper and brown basal layers, overlying white calcareous shell. O, *Mutela legumen*, showing two periostracal layers: upper yellow layer and blue basal layer, in addition to blue coloured calcareous shell matrix. Coloured arrows indicate the location of colour: black-filled circles indicate organic material (periostracum/organic layers in calcareous shell), black outlined circles indicate inorganic material (calcareous shell matrix). Raman measurements were taken in all circled locations. Black lines indicate periostracum. White arrows indicate calcareous shell. Species figured in the same order as in Figure 1.

### Raman spectroscopy

Using the cross-sectional images as a guide, Raman measurements were taken from all coloured areas, with a focus on green and blue regions of colour in the periostracum and from calcareous shell (see Fig. 2 for location of Raman measurements).

Measurements were taken using a Renishaw Virsa instrument at the Natural History Museum with 532 and 785 nm excitation in back-scatter configuration with a 50X objective lens linked by fibre optic cables to the spectrometer, producing a laser spot of ~5 μm diameter. Measurements using 532 nm excitation were acquired using a laser power of 0.1 mW (set using a variable angle shutter) and were summed over 20 accumulations of 0.1 seconds. Measurements using 785 nm excitation were acquired using a laser power of 5 mW and summed over 5 accumulations of 10 seconds. Exposure times were adjusted on a sample-by-sample basis to avoid signal saturation due to intense sample fluorescence. A 2,400 lines/mm grating was used to produce a spectral range of 100–3200 cm^−1^ measured over multiple acquisitions in stepped mode, producing a spectral bin size of ~1.3 cm^−1^ (532 nm) and ~0.9 cm^−1^ (785 nm) per pixel. Cosmic rays were removed using the Renishaw WiRE software, all further Raman data processing (background subtraction, peak detection, peak fitting, data visualisation) was done using a Jupyter Notebook running custom Python code (21) that utilises the following packages: Numpy (22), SciPy (23, 24), and LMFIT (25). Initial background subtraction was done by fitting a polynomial function to regions with no Raman signal and subtracting it from the data. A second background subtraction was then done to reduce the magnitude of background oscillations by fitting and subtracting a background reference spectrum, averaged from 9 (532 nm) and 11 (785 nm) spectra that exhibited a maximum absolute first derivative of <0.021 between 200 and 1800 cm^−1^ (see Supplementary Figure S2). An automatic peak detection algorithm was used to identify local maxima with signal:noise ratios exceeding 10:1 and a relative intensity higher than 5% vs the maximum observed signal. Automatic peak fitting was done using pseudo-Voigt functions to represent each detected Raman peak. The Jupyter Notebook and underlying code are available on GitHub and archived at (21).

## Results

### Optical analysis

In every shell with external green colouration (Figs. 1A-D, J-N), a single green layer was observed in the middle or basal layer of the periostracum (closest to the calcareous shell), but never in the upper layer (Table 1, Figs. 2A-D, J-N). Green colour was entirely absent from the calcareous shell matrix.

**Table 1.**
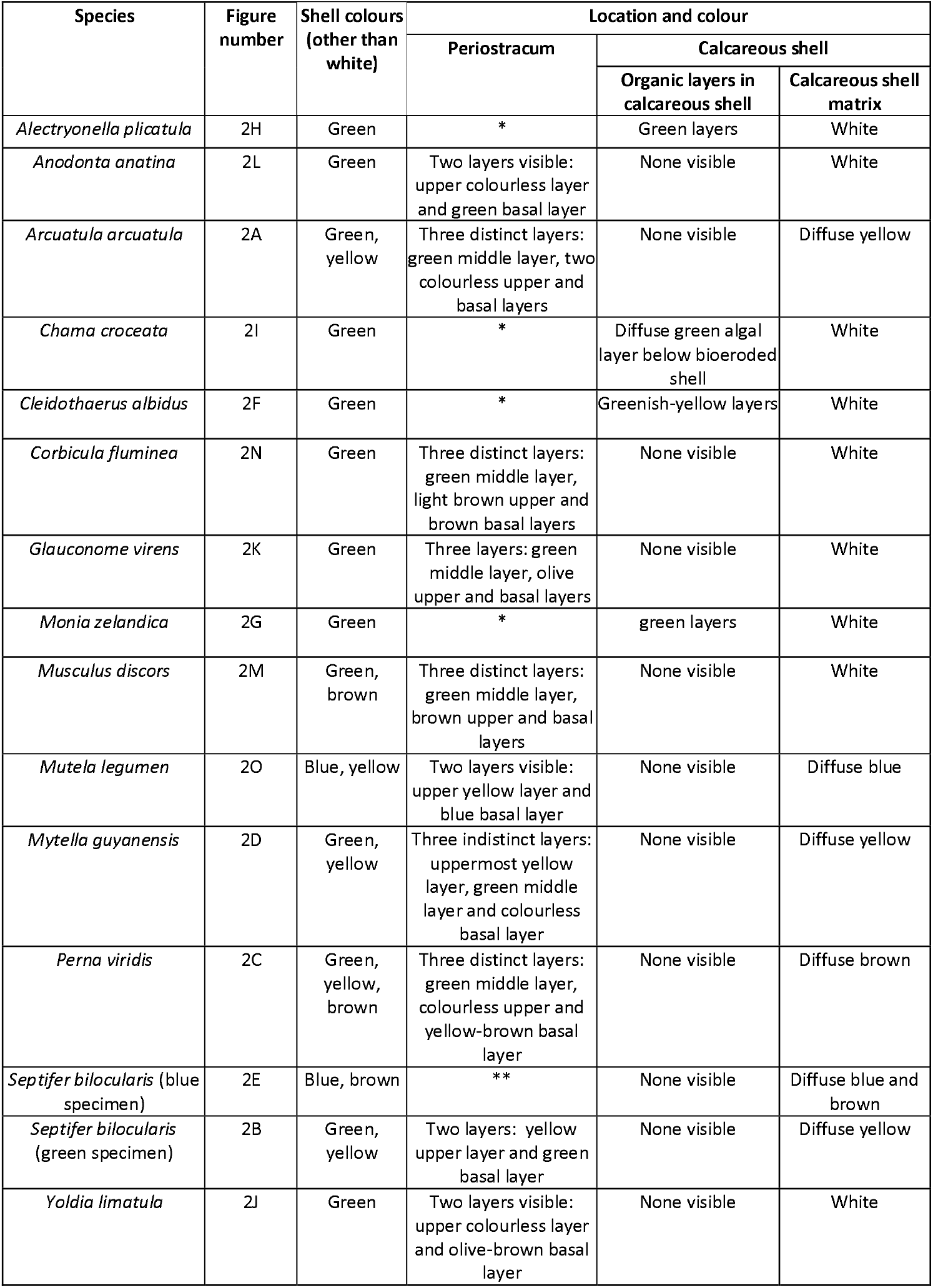
Summary observations of shell colour location and optical microscopy results linked to Figure 2. * periostracum ≤ 1µm; ** periostracum removed from the specimen prior to purchase.

Almost all species with internal green shell colouration displayed multiple green organic layers interleaved between calcareous shell layers (Table 1, Figs. 1F-I and 2F-I). A greater number of green layers corresponded with a deeper visible shell colour.

*Chama croceata* differed from all other shells with internal green colouration in that discrete organic layers were not present, either in the periostracum or calcareous shell. Instead, a green region was identified in the base of the outer calcareous shell, which could be observed from inside the shell, in addition to green pigmented diatoms which were embedded in the heavily bioeroded outermost region of the shell (see Fig. 2I and inset).

In some of the shells with green periostracal colour, an additional diffuse yellow (Figs. 2A - *A. arcuatula*, B – *Septifer bilocularis*, D – *Mytella guyanensis*) or brown area (Fig.2C – *Perna viridis*) was observed in the underlying calcareous shell matrix.

The two species with external blue shell colour, *S. bilocularis* and *Mutela legumen*, had areas of diffuse blue colour in the calcareous shell matrix (Fig. 2E and O, respectively). In the case of *S. bilocularis*, which was missing its periostracum, there were additional brown streaks in the shell (Fig. 2E). Blue and brown areas corresponded to externally visible areas of the same colour (see Figs. 1E and 2E). In *M. legumen* an additional single distinct blue layer in the base of the periostracum was also present (Table 1, Fig. 2O).

### Raman analysis

#### Green

In contrast to other colours, Raman spectra for all organic green layers were very noisy and no clear carotenoid-based pigment peaks could be identified at either 532nm or 785nm excitation wavelengths (Figs. 3A and 3B respectively, see S3 for non-normalised spectra, Fig. 4 for SNR graphs).

**Figure 3.**
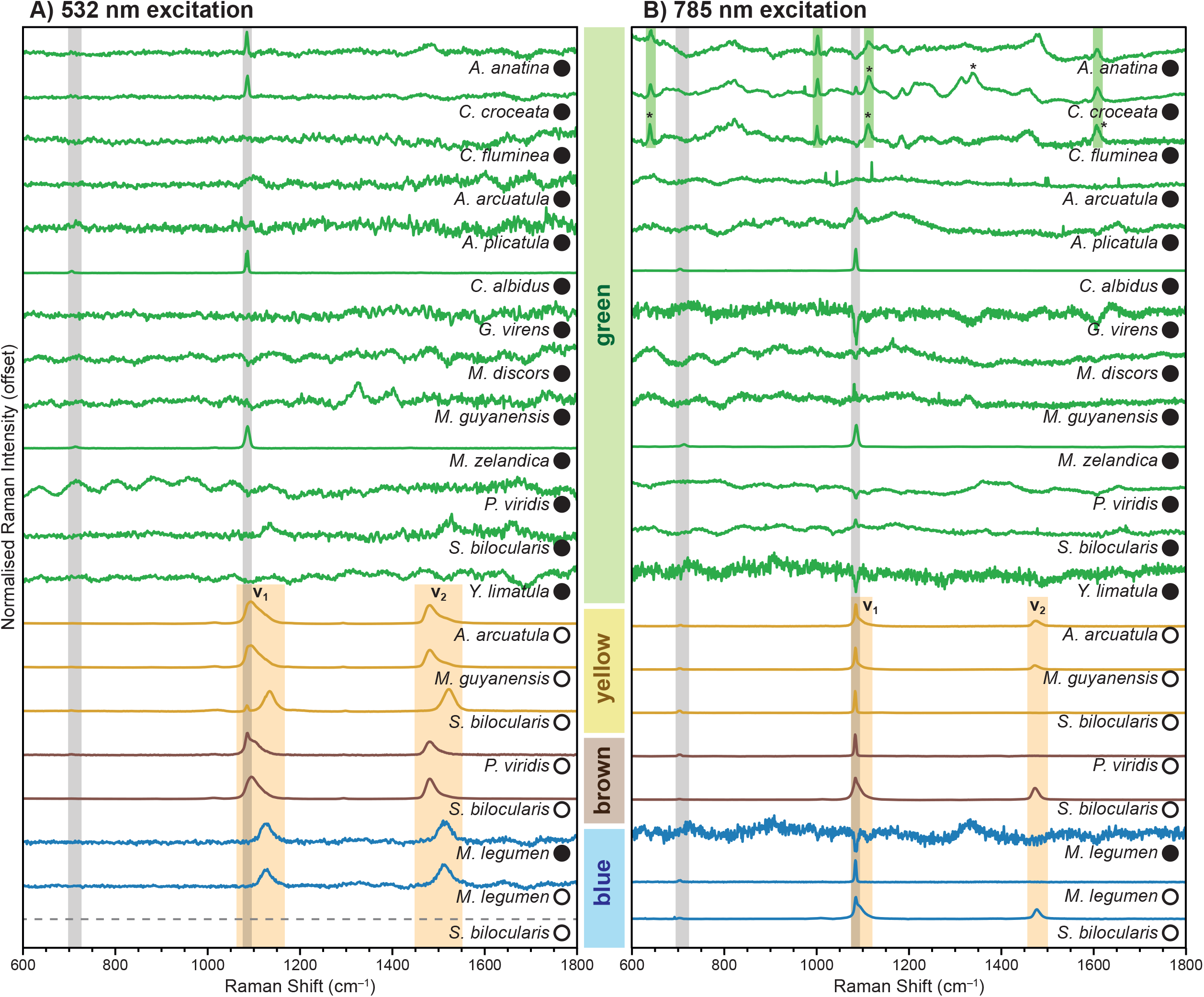
Raman spectra of 15 green or blue coloured bivalve species, acquired using (A) 532 nm or (B) 785 nm excitation. Spectra were baselined, background oscillation subtracted, normalised and offset for clarity. Solid circles indicate measurements taken of organic regions (periostracum/organic layers in calcareous shell); open circles indicate inorganic shell matrix. Spectrum colours reflect shell colour, samples are grouped according to apparent colour (green, yellow, brown, blue). Vertical grey bars indicates the position of the calcium carbonate peak, yellow bars indicate key carotenoid-based peaks, green bars in the 785 nm spectra mark potential peaks of unknown pigment. Asterisks indicate all significant non-carotenoid-based peaks identified using autodetection allowing for a signal:noise ratio of 10:1. No data were collected for *S. bilocularis* (blue) using 532 nm excitation due to machine failure.

**Figure 4.**
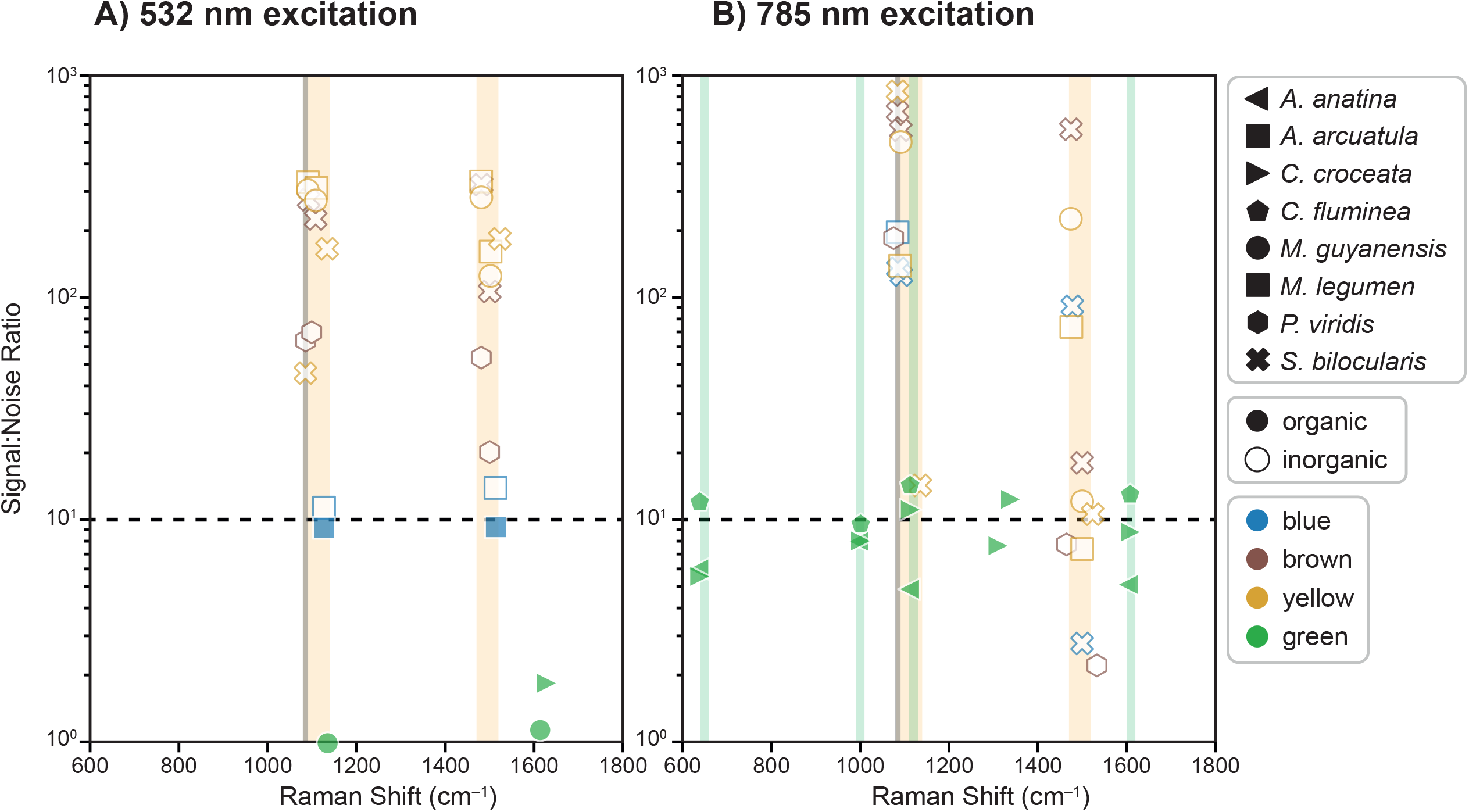
Signal:noise ratio (SNR) of Raman pigment peaks observed in eight species at (A) 532 nm and (B) 785 nm excitation. Peaks were identified automatically using pseudo-Voigt function to represent each detected Raman peak. Noise was calculated as the standard deviation of the peak-free region 1700–1800 cm-1. Solid symbols indicate measurements taken of organic regions (periostracum/organic layers in calcareous shell), open symbols indicate measurements of inorganic calcareous shell matrix. Vertical grey bar indicates the expected position of the calcium carbonate peak, vertical orange bars indicate the expected position of carotenoid-based peaks, vertical green bars indicate the apparent positions of peaks associated with an unknown pigment with weak Raman activity using the 785 nm excitation. Colour of symbol reflects colour of shell. Horizontal black dashed line indicates the 10:1 SNR threshold required for unambiguous positive peak detection. Seven species with no significant pigment peaks in green periostracum were excluded from the graph.

**Figure 5.**
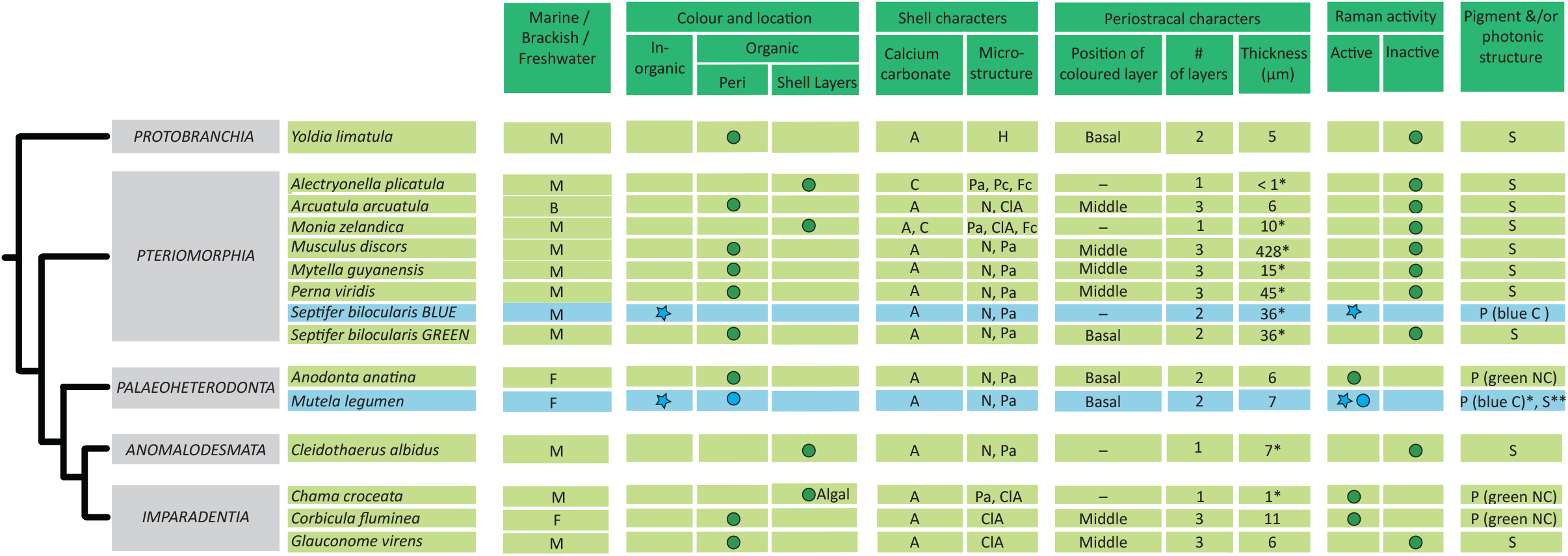
Simplified phylogeny of Recent bivalves (modified from Bieler et al., 2014) showing mapped traits. *Aquatic environment* (from (45, 46)): M – Marine, B – Brackish, F – Freshwater; *Colour and location*: Inorganic - calcareous shell matrix, Organic peri – periostracal layers, Organic shell layers – organic layers interleaved between calcareous shell; *Shell characters*: Calcium carbonate crystal: A – Aragonite, C – Calcite, ClA – crossed lamellar aragonite, Fc – Foliated calcite, H – Homogeneous (no layers), Pa – Prismatic aragonite, Pc – Prismatic calcite, N – Nacreous aragonite. Information from various sources, including (47–49). *Periostracal characters*: Periostracal thickness values, * from Harper (50). All other values were measured by the authors from scanning electron micrographs. Suggested mechanism of shell colour based on results of this study: P – Pigment, S – Photonic structure, * - blue coloured calcareous shell matrix is due to a carotenoid-based pigment, ** blue periostracal colour could be due to a photonic structure.

Novel non-carotenoid-based peaks were identified in three species, *C. fluminea, A. anatina* and *C. croceata*, and observed using 785nm excitation only (Fig. 3B). Four peaks identified in these species were shared (650 cm^−1^, 1002 cm^−1^, 1115 cm^−1^ and 1609 cm^−1^; locations of these peaks are highlighted with green bars in Fig. 3B). Several of the observed non-carotenoid peaks are weak and difficult to consider unambiguous detections of real Raman-scattering phenomena, with signal:noise ratios (SNRs) of less than 10:1 (Fig.4). *Corbicula fluminea* had three peaks that exceeded the conservative 10:1 threshold for unambiguous detection, at 650 cm^−1^, 1115 cm^−1^ and 1609 cm^−1^ (indicated by asterisks in Fig. 3B), while the 1002 cm^−1^ peak was close to the threshold (Fig. 4A). An unambiguous peak was also detected at 1115 cm^−1^ in *C. croceata* (indicated by an asterisk in Fig. 3B). In *C. croceata* only, a further two unique pigment peaks were observed at 1312 and 1340 cm^−1^, of which only the 1340 cm^−1^ peak was unambiguous (indicated by an asterisk in Fig. 3 B).

#### Blue

Of the two blue species examined, two identical peaks corresponding to carotenoid-based pigments were observed at 1128 and 1514 cm^−1^ in the organic periostracum and calcareous shell matrix of *M. legumen* under 532 nm excitation (Fig. 3A). Only the shell matrix peaks were unambiguous, with SNRs around 11:1 (Fig. 4B).

In the second species, *S. bilocularis*, carotenoid-based pigments were also detected, but these differed from those found in *M. legumen*, with different peaks at 1090 and 1478 cm^−1^ in the blue calcareous shell matrix using 785 nm excitation (Fig. 3B). These peaks were unambiguous with SNRs of approximately 100:1 (Fig. 4A).

#### Non-rare colours (brown and yellow)

Additional yellow and brown calcareous shell matrix colours demonstrated carotenoid-based pigment peaks (Figs. 3A and B). Under 532 nm excitation, two major peaks (*v*_1_, *v*_2_) associated with yellow carotenoid-based pigments were observed at 1091 cm^−1^ and 1481 cm^−1^ in *A. arcuatula*, at 1090 cm^−1^ and 1481 cm^−1^ in *M. guyanensis*, and at 1134 cm^−1^ and 1522 cm^−1^ in green *S. bilocularis* (Fig. 3A). These peaks were also observed under 785 nm with more consistent positions of 1093-1094 cm^−1^ and 1475-1476 cm^−1^ (Fig. 3B). Two minor carotenoid-based pigment peaks (v_3_: at ~1010 cm^−1^, v_4_: ~1300 cm^−1^) were only observed under 532 nm excitation (Fig. 3A). All peaks were unambiguously detected using this excitation, with SNRs between 11:1 and 400:1 (Fig. 4B).

Carotenoid-based pigment peaks for the green and blue *Septifer bilocularis* shells were non-identical.

For brown calcareous shell matrix colour, *S. bilocularis* showed carotenoid-based pigment peaks at 1095 cm^−1^ and 1482 cm^−1^ (versus 1090 and 1478 cm^−1^ in blue shell). *Perna viridis* showed peaks at 1086 cm^−1^ and 1482 cm^−1^. All peaks were unambiguous under 532nm excitation, with SNRs between 50:1 and 100:1 (Fig. 4B).

As in previous studies, carotenoids were easier to detect using 532 nm excitation than 785 nm, the former producing strong Raman peaks that were non-symmetric in shape and varied significantly in position (~43 cm^−1^) between species (suggesting non-identical pigments) due to resonance enhancement of vibrations associated with the carotenoid chromophore at this excitation wavelength (Fig. 3A). By comparison, under non-resonant 785 nm excitation the carotenoid peaks were harder to detect, more symmetric, and showed less variation in positions between species (<5 cm^−1^) (Fig. 3B).

#### Calcium carbonate

The calcium carbonate matrix of the calcareous shell was also apparent in Raman spectra of several species, having peak positions consistent with either calcite (280, 712, 1085 cm^−1^) or aragonite (208, 704, 1085 cm^−1^) (Figs. 3A and B). Calcite was only detected in *A. plicatula* and *M. zelandica*; whereas aragonite was detected in *A. anatina, A. arcuatula, C. croceata, C. albidus, C. fluminea, G. virens, M. legumen, M. guyanensis, P. viridis, S. bilocularis* (green & blue) *and Y. limatula*. In *M. laevigata*, only the 1085 cm^−1^ peak common to both calcite and aragonite was observed.

#### Trait mapping

Mapping traits onto a simplified bivalve phylogeny revealed that non-carotenoid-based green pigments occur only in the Palaeoheterodonta and Imparadentia, and within these clades, only in freshwater species. *Chama croceata* is the exception since it is a marine species, and colour is likely due to algae in the calcareous shell.

Pigments were not detected from any green Pteriomorphia specimens examined, nor any species of Protobranchia or Anomalodesmata. All species examined in this study from these clades are marine, except *Arcuatula arcuatula*, which occurs in brackish water.

Two different carotenoid-based blue pigments were identified in two species, one each from the Pteriomorphia and Palaeoheterodonta.

There does not appear to be any correlation between colour mechanism and shell microstructure, apart from to note that in the two clades in which blue was found, both times it was associated with nacre and prisms.

## Discussion

Bivalve shells are often brightly coloured, with all the major colour groups represented. Here we have investigated the mechanisms by which the rarest of colours, blue and green, are produced in a phylogenetically diverse group of bivalve species. We present results from both Raman spectroscopy and general observations and reveal that these colours are produced by multiple, independent pathways.

### Rare colours: Green

Green colour across all clades was observed only in organic parts of the shell, either as distinct layers in the periostracum or in organic layers within the calcareous shell, supporting the idea proposed by Grant and Williams (3), that rare green shell colour occurs more often in organic, than inorganic material.

No significant carotenoid-based pigment peaks were detected in any green-coloured shell. This result was also reported by Wade et al., (17) for a smaller number of samples. However, non-carotenoid-based peaks that may be associated with a pigment were identified in three species (*C. fluminea, A. anatina* and *C. croceata*). Four peaks observed in *C. fluminea* exceeded the 10:1 signal:noise ratio (SNR) required for unambiguous detection against background noise. These same peaks were also found in *A. anatina* and *C. croceata*, albeit not all peaks in these species meet our conservative SNR threshold of 10:1. The fact that these four peaks were shared across three species, with similar positions and widths, suggests, however, that they are non-trivial and are not the result of random fluctuations or sample contamination by other materials. The observed peak positions may indicate the presence of an aromatic chromophore on the molecule and therefore suggests it may have a role in pigmentation (26). However, the observed positions are not consistent with carotenoids (the most common class of pigment in bivalve shells), nor are they consistent with melanins or tetrapyrroles, which also occur in mollusc shells (27–30). The existence of novel pigments with weak Raman activity was first suggested by Wade et al. (17), but this is the first direct detection of such pigments in mollusc shells, and as such they therefore warrant further investigation.

The absence of significant pigment peaks in the remaining ten species with green-coloured shells could mean that there may be pigments that are Raman active but present at such low concentrations that they remain undetectable to the measurement settings used in this study. However, this is doubtful, since both this study and Wade et al.’s (17) used resonant excitation wavelengths that are resonant with most visible pigmentation, enhancing sensitivity to such compounds even at very low concentrations. The lack of detectable pigment peaks in these species suggests the presence of structural colour. Our results provide support for the more widespread existence of structural colour in bivalves, which has until now been reported only in the pteriomorph species, *Perna canaliculus* and *P. viridis* (18, 19). Both species possess matt green externally coloured shells, but the middle layer of the periostracum appears iridescent green. Pan et al. (19) showed *P. canaliculus* has obliquely arranged nanofibres, which act as a photonic structure to diffract light and generate the green iridescence. However, the authors were unable to explain how the green iridescence contributes to the matt green external shell colour, since the iridescence was humidity-dependent and disappears when dry, but the external green shell colour persisted. This led them to conclude that the external green colour must be due to pigments. Our results show that there are no detectable pigments in the green periostracal layer of *P. viridis*, which would suggest the green shell colour is due to the photonic structure. In fact, the green periostracal layer did not appear iridescent in our study (Fig. 2), possibly because the resin-embedding process filled some of the air spaces in the periostracum. We did, however, identify a brown carotenoid-based pigment in the calcareous shell matrix, which would provide a contrasting background to the structural green iridescence of the periostracum. Structural and pigmentary elements have been found working in concert in almost all other taxa in which structural colour has been described (31).

*Chama croceata* was the only species examined to host algae in the shell. Two different types were identified from different shell regions. Diatoms (possibly from the genus *Navicula*) were recorded from recesses within the bioeroded shell, similar to those reported from the shell surface of other bivalves (32), although not *Chama* specifically and with no reference to the alteration of overall shell colour. In addition to diatoms, an indistinct green layer was observed, thought to be a filamentous endolithic green alga, similar to *Ostreobium* reported from, and contributing to the green colour of some corals (33). The green of both types of algal cells seen against the white shell of *C. croceata* appear to be the source of green shell colour, though the weak Raman signal of the novel, non-carotenoid based pigment did not correspond to known spectra for chlorophyll (e.g.(29)).

### Rare colours: Blue

Unlike green, blue was observed both as a layer in the organic periostracum but also as a diffuse region in the calcareous shell matrix in the two species examined. Significant carotenoid-based pigment peaks were detected in both blue species, *M. legumen* and *S. bilocularis*, from calcareous shell with diffuse blue colour. Carotenoids can only be synthesised by plants and fungi and are therefore almost always acquired by animals, including bivalves, through their diet (34). Plant carotenoids are normally yellow, orange and red, so we speculate that the pigment has been chemically or structurally modified with a protein in *M. legumen* and the blue *S. bilocularis* to produce a blue pigment, as has been reported previously in other invertebrates (35, 36).

The spectra from both periostracum and calcareous shell in *M. legumen* appeared to be identical, but the signal was non-significant from the periostracum. While this may reflect the presence of pigments in the periostracum, another hypothesis is that the similarity might be explained by the proximity/overlap of the laser spot across periostracum and shell, since the spot size was of a similar diameter (5 µm) to the periostracal thickness (6 µm), and the spot may be effectively enlarged by scattering of laser light within the solid matrix of the sample itself (37). As a result, the strong carotenoid response from the shell would be detected even if the laser spot was centred on the periostracum. If the periostracum spectrum is due to an ‘overlap’ with pigments found in the calcareous shell, the blue periostracal colour of *M. legumen* may be structural in origin. However, we did not find evidence for ‘overlap’ readings in another species with a thin periostracum and nearby carotenoid-based pigment (for example, see *A. arcuatula* spectrum in Figure 4b), which would suggest that *M. legumen* spectra correctly reflect the existence of a blue pigment in both periostracum and shell in this species.

### Evolution of traits

General observations of 13 green bivalve shells showed that green-coloured shells are not only most common in the Pteriomorphia and Palaeoheterodonta, but these clades also include the most vividly green-coloured specimens. Species with blue-coloured shells are most common in Palaeoheterodonta (3) but vividly coloured specimens occur in both Palaeoheterodonta and Pteriomorphia.

Mapping colour-related traits for these rare colours onto a simplified bivalve phylogeny revealed that non-carotenoid-based green pigments occur only in the Palaeoheterodonta and Imparadentia (given limited sampling to date). Two of the three species examined occur in freshwater, excluding *Chama croceata*, which is marine with algae in the calcareous shell. In contrast, green species from the Pteriomorphia, Protobranchia and Anomalodesmata, without detectable pigments and thought instead to possess photonic structures - are all marine species, except *Arcuatula arcuatula*, which occurs in brackish water. Taken together, these results demonstrate that within Bivalvia, clades have convergently evolved green shell colour via two different mechanisms, providing further evidence for Grant and Williams’s (3) idea that there is likely to be a selective advantage for green bivalve shell colour.

Blue-shelled species are rare, but in the two examined, two different carotenoid-based pigments are thought to be the cause of colouration, with blue colour found in the periostracum and the calcareous shell matrix, although the possibility that periostracal colour may be structural cannot yet be entirely ruled out. It is difficult to draw conclusions based on such a small sample size, but the results could suggest that blue is more commonly produced by pigments rather than structures. This is unusual compared to other taxa, where the colour blue is more often structural, for example, in blue gastropods (38) and cephalopods (39), and also in unrelated taxa including, blue birds (40) and butterflies (41)).

### Future work

Future work will aim to identify and characterise photonic structures in green pteriomorphs, where the most vivid colours and strongest evidence of these structures are found. The structural green colour in this group has been linked to the presence of a vacuolar layer in the periostracum (18, 19) and its evolution may be driven by a multitude of factors, including the need for increased mechanical strength in habitats with greater wave exposure (42). This periostracal layer is absent in the freshwater palaeoheterodont clade (for example, (43, 44)), which may provide further clues about its evolution. We speculate that structural green colour has evolved in pteriomorphs through changes in the size of the vacuoles in the vacuolar layer. The resultant colour may provide a selective advantage by means of camouflage and a reduction in predation. Based on these hypotheses, we make the following predictions: 1) Green pteriomorphs will have vacuolar layers with vacuoles appropriately sized to reflect green light. 2) Green pteriomorphs with a vacuolar layer will be found in the top 200 m of the ocean (the photic zone), where the colour will be visible to potential predators. These predictions require testing and are the focus of ongoing studies.

## Acknowledgements

We would like to thank Dale Boorman at Renishaw and Wren Montgomery at the Natural History Museum for organising and providing technical support for the loan of the Virsa Raman instrument used in this study. Chris Howard, Yidan Wang and Donatella Banti are thanked for Raman advice and preliminary data collection. Tom White provided access to specimens and Aimee McArdle, Kevin Webb, Lucie Goodayle and Jonathan Jackson took all photographs. Callum Hatch is thanked for slide preparation of all shells. Dana Perry provided Hirox training and guidance. John Taylor and Liz Harper are thanked for helpful discussion about bivalves. Martin Cheng and Kerry Walton are thanked for specimen collection. Eileen Cox is thanked for diatom identification.

## Supplementary Information

**Figure S1**. All 15 blue and green bivalve mollusc species examined for the study, photographed against the Munsell Colour Chart. Images were taken with a QPcard 101 to allow for colour correction of final images.

**Figure S2**. Illustration of Raman spectrum processing steps: 1) polynomial baseline subtraction to remove major background features. 2) fitted reference subtraction to reduce the impact of the background oscillation. 3) automatic peak detection based on an SNR threshold of 10:1 and a minimum relative intensity of 5%. 4) automatic peak fitting using one pseudo-Voigt function per detected peak with a window of ±150 cm^−1^ to estimate peak positions, full-width-half-maxima (FWHM), and SNRs.

**Figure S3**. Raman spectra of 15 green or blue coloured bivalve species, acquired using (A) 532 nm or (B) 785 nm excitation. Spectra were baselined, background oscillation subtracted and offset for clarity. Solid circles indicate measurements taken of organic regions (periostracum/organic layers in calcareous shell), open circles indicate inorganic calcareous shell matrix. Spectrum colours reflect shell colour, samples are grouped according to apparent colour (green, yellow, brown, blue). Vertical grey bars indicate the position of the calcium carbonate peak, yellow bars indicate key carotenoid-based peaks, green bars in the 785 nm spectra mark potential peaks of unknown pigment. Asterisks indicate all significant non-carotenoid-based peaks identified using autodetection allowing for a signal:noise ratio of 10:1. No data were collected for *S. bilocularis* (blue) using 532 nm excitation.

