## Supplementary figures and images for "Rare shell colours in bivalves reveal multiple evolutionary pathways to blue and green colouration"

### Supplemental Figure 1

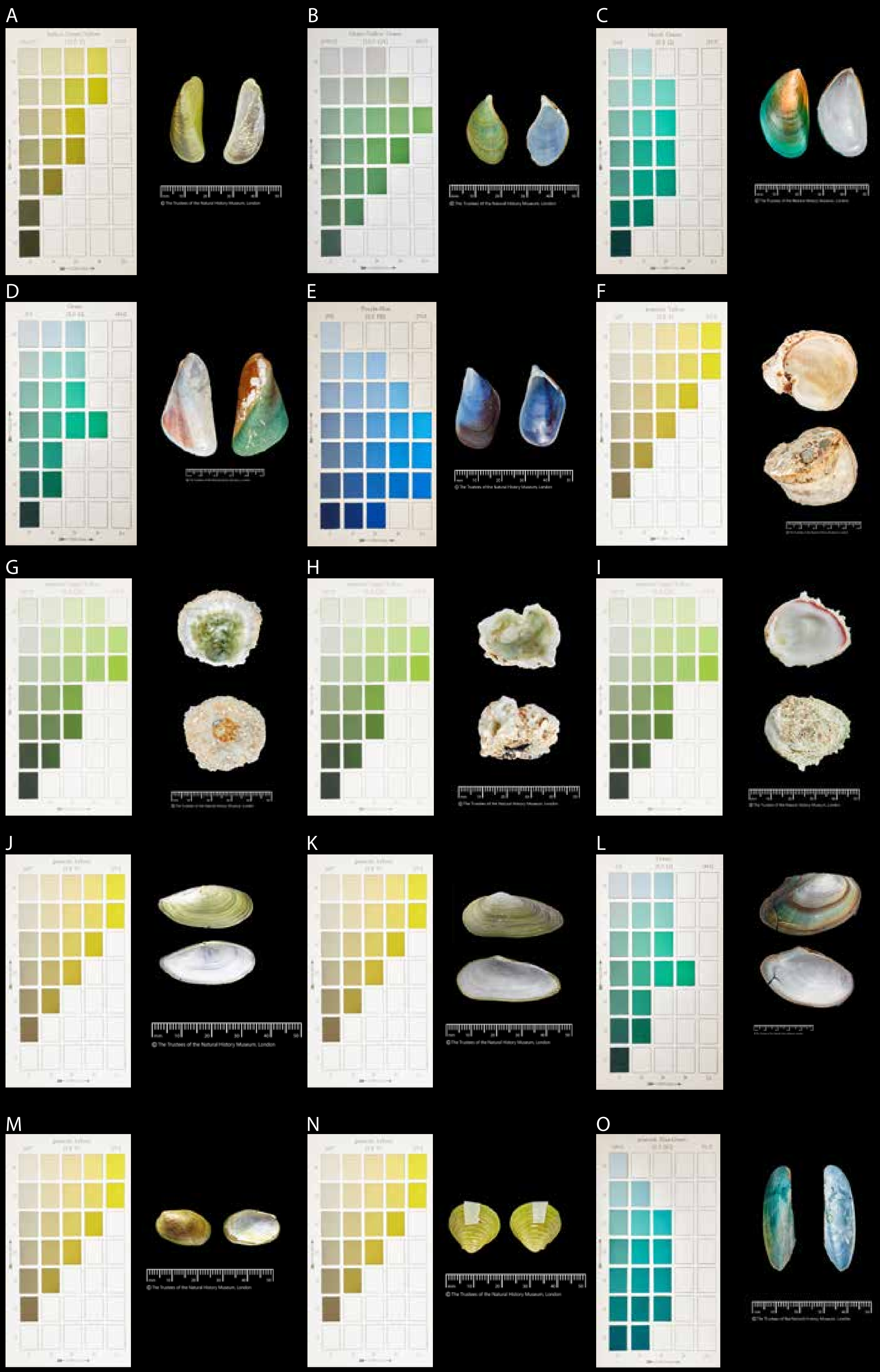

### Supplemental Figure 2

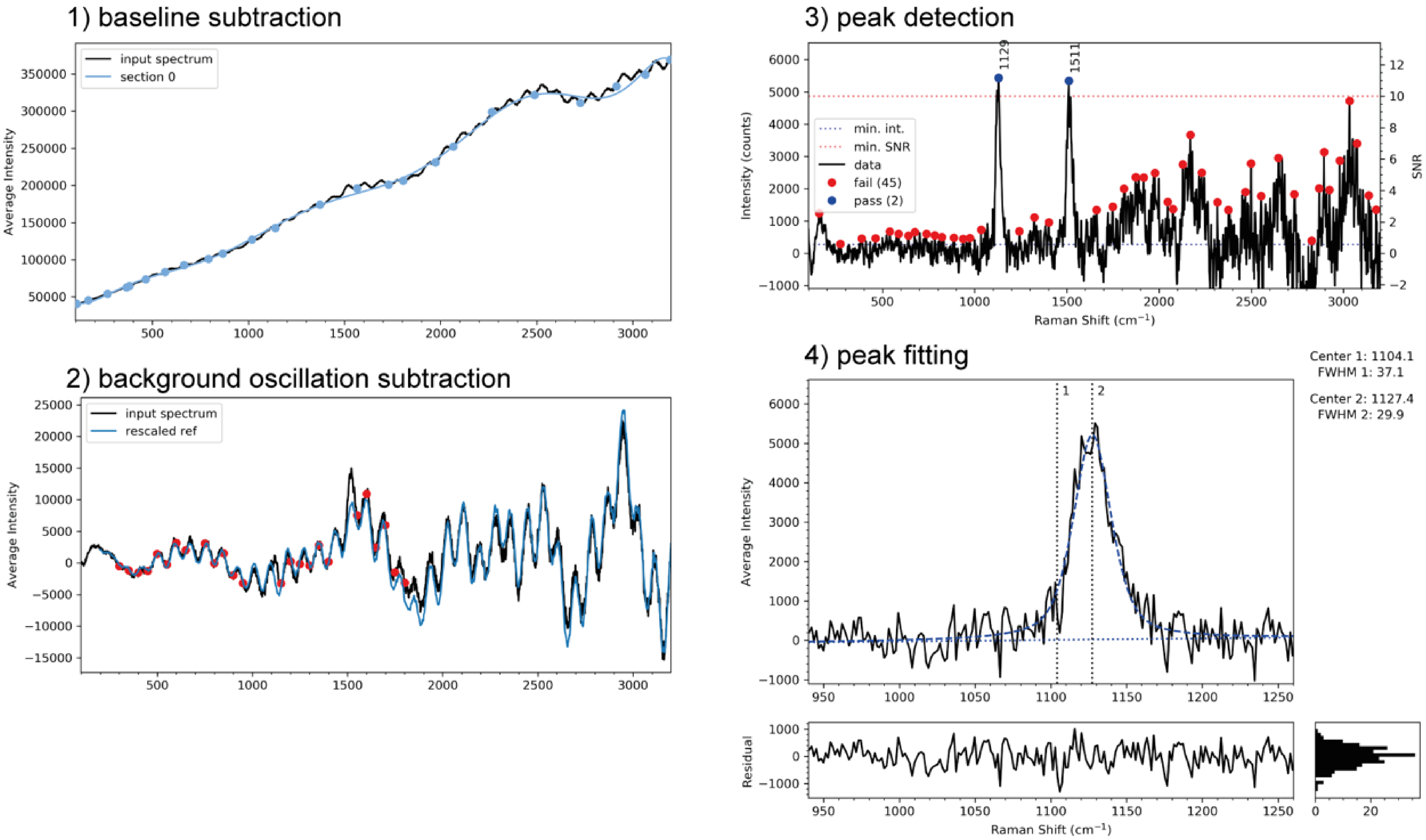

### Supplemental Figure 3

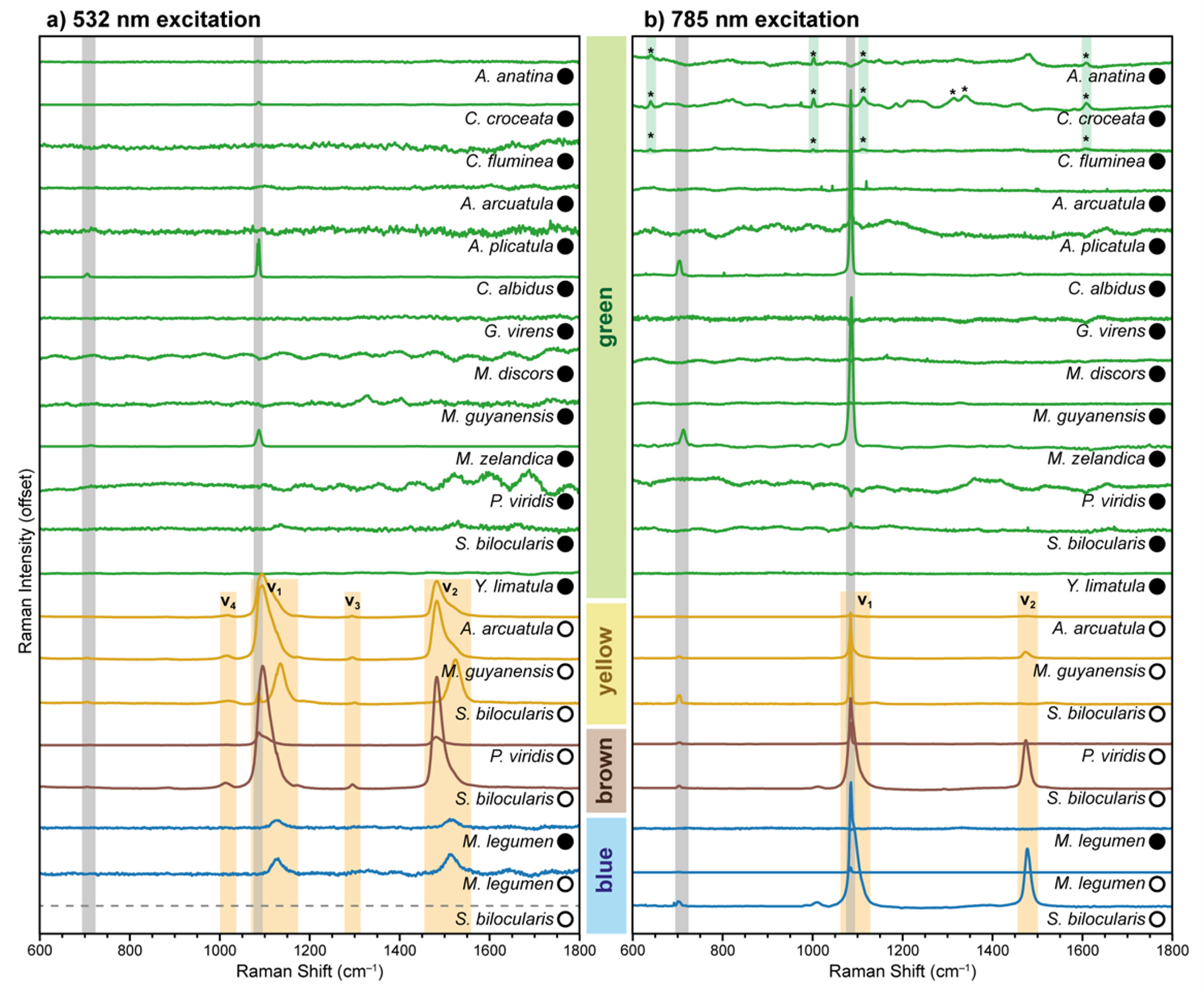
